# LEF1 isoforms regulate cellular senescence and aging

**DOI:** 10.1101/2023.07.20.549883

**Authors:** Minxue Jia, Khaled Sayed, Maria G. Kapetanaki, William Dion, Lorena Rosas, Saad Irfan, Eleanor Valenzi, Ana L. Mora, Robert A. Lafyatis, Mauricio Rojas, Bokai Zhu, Panayiotis V. Benos

## Abstract

**Background:** The study of aging and its mechanisms, such as cellular senescence, has provided valuable insights into age-related pathologies, thus contributing to their prevention and treatment. The current abundance of high throughput data combined with the surge of robust analysis algorithms has facilitated novel ways of identifying underlying pathways that may drive these pathologies.

**Methods:** With the focus on identifying key regulators of lung aging, we performed comparative analyses of transcriptional profiles of aged versus young human subjects and mice, focusing on the common age-related changes in the transcriptional regulation in lung macrophages, T cells, and B immune cells. Importantly, we validated our findings in cell culture assays and human lung samples.

**Results:** We identified Lymphoid Enhancer Binding Factor 1 (LEF1) as an important age-associated regulator of gene expression in all three cell types across different tissues and species. Follow-up experiments showed that the differential expression of long and short LEF1 isoforms is a key regulatory mechanism of cellular senescence. Further examination of lung tissue from patients with Idiopathic Pulmonary Fibrosis (IPF), an age-related disease with strong ties to cellular senescence, we demonstrated a stark dysregulation of LEF1.

**Conclusions:** Collectively, our results suggest that the LEF1 is a key factor of aging, and its differential regulation is associated with human and murine cellular senescence.

## BACKGROUND

It is generally accepted that as the world’s population is getting older due to the increase in life expectancy^1^, the need of preventing and addressing age-associated pathologies will increase as well. Therefore, it is necessary to heighten our effort in understanding the complex mechanisms underlying healthy aging as opposed to disease-ridden survival. On the molecular and cellular level there are 12 identified hallmarks of aging^2^, which form a framework for age-related research. Cellular senescence, identified as one of the hallmarks of aging, is contributing to aging either through the accumulation of damaged senescent cells or the secretion of pro-inflammatory cytokines and matrix metalloproteases, often referred to as Senescence Associated Secretory Phenotype (SASP)^3,4^. Cumulative evidence supports the existence of rather ubiquitous age-associated molecular mechanisms which can occur in multiple tissues/organs of the same organism or across different species^5^. Research on multiple organs and organisms suggests that addressing the underlying mechanisms of aging could impact the course of several pathologies that are currently associated with old age.

While common mechanisms may underlie the process of aging in different organs, the unique physiology of each one of them as well as environmental stressors will also determine how aging impacts their fitness. The lung, as an organ, presents a unique system due to its cellular complexity of at least 40 discrete cell types^6^ with a large interface surface that is constantly challenged by diverse stressors. Advanced age is a significant risk factor for several lung diseases including COPD^7^ and IPF^8^ which interestingly present some of the same irregularities in cellular mechanisms that are considered hallmarks of aging, including cellular senescence^9^. Due to these similarities, the investigation of age-related molecular changes could lead to a better understanding of disease pathogenesis and eventually more efficient therapeutic interventions.

In recent years, the abundance of molecular high throughput data has supported the development of a plethora of functional and mechanistic computational analysis tools which are widely used to analyze gene expression data. Importantly, many of these transcription factor (TF) and pathway analysis methods are successfully adapted for single cell RNA-seq (scRNA-seq) data^10^ and can provide meaningful insights into single cell biology.

In the current work, we analyzed scRNA-seq and bulk RNA-seq data from human lung and blood samples of healthy donors as well as mouse lung samples, focusing on macrophages, T cells, and B cells which were abundant and well-identified in all datasets. Interestingly, we didn’t find any differentially expressed genes (young *vs*. aged) shared between human and murine immune cells. However, when we examined the transcription factor activity (regulons), we were able to identify Lymphoid Enhancer-binding Factor 1 (LEF1) regulon as a common mechanism of aging in human and murine lungs and in human blood. More importantly, we provide experimental evidence to support a direct functional role of the long LEF1 isoform in partially reversing cellular senescence through a possible new regulatory mechanism.

## MATERIALS AND METHODS

### Datasets

For this project we used four publicly available datasets. **(1)** <u>GSE128033</u>, which includes human scRNA-seq samples taken from whole lung tissue of three young (age ≤23, 14,704 cells), two heathy aged (age ≥55, 8,573 cells), and six Idiopathic Pulmonary Fibrosis (IPF) donors (age ≥69, 26,538 cells). This dataset was generated by our group^11,12^. (2) <u>GSE122960</u>, which includes human scRNA-seq samples of whole lung tissues of two young (15,770 cells) and three aged (21,981 cells) healthy donors^13^. **(3)** <u>GSE124872</u>, which includes mouse scRNA-seq samples from whole lung tissue of seven 3-month-old (7,672 cells) and eight 24-month-old mice (7,141 cells)^14^. **(4)** <u>GSE158699</u>, which includes bulk RNA-seq data from blood samples from former and current smokers of the COPDGene study^15^. This dataset was divided in two groups: young (n = 79, age ≤55) and aged (n = 123, age ≥75).

For human scRNA-seq data, we used all samples (never-smokers and former smokers; n=10, 60,922 cells) to identify main cell types via unsupervised clustering (package: Seurat^16^). Subsequently, we took special care to select healthy samples that are free of smoking-related or other pathologies by carefully examining the provided records of smoking status and lung histopathology. We did so to ensure that the identified signals can be attributed to aging and not any aging confounders. This process eliminated other publicly available datasets we considered and resulted in the exclusion of two samples from the above datasets. Donor 4 was a former smoker with abnormal cell type distribution (Figure S2); and Donor 2 was excluded due to the histopathology that indicated previous smoking or pollution exposure (Figure S3). Complete list of samples used in cell type identification and presented in Table S1.

### Digital cytometry

We used CIBERSORTx^17^ to infer cell-type proportions and cell-specific gene expression from the blood-derived bulk RNA-seq data. Deconvoluted cells with an imputed number of genes <5% of the total number in the bulk RNA-seq dataset were excluded from our analysis. Only CD4^+^ Naïve T cells had an imputed number of genes >5%.

### Differential Gene Expression data analysis

We used Seurat V3^16^ on the human and mouse scRNA-seq data to determine differential gene expression between aged and young lungs. For the Mouse Lung dataset (ML), we first created a Seurat object for the raw counts using CreateSeuratObject. Then, we calculated the percentage of the mitochondria genes (percent.mt; function: PercentageFeatureSet). Cells with >5% mitochondrial genes (variable: percent.mt) and with number of features <250 or >2000 (variable: nFeatures_RNA) were filtered out. The filtered data were then normalized and scaled (functions: NormalizeData and ScaleData). We also used FindVariableFeatures to find the top 2000 high variable genes. For the Human Lung dataset (HL), we filtered out cells with high percentage of mitochondria genes (percent.mt >35%) and with number of features less than 200 (nFeatures_RNA). Besides, we filtered out empty droplet and doublets identified by Scrublet and emptyDrops^18,19^. The filtered (missing) data were then imputed using SAVER^20^. The imputed data were normalized and scaled using the sctransform function, and the number of UMIs per cell as well as the percentage of mitochondrial gene content were regressed out.

In order to create clusters for the different cell types, we performed Principal Component Analysis (PCA) to reduce the dimensionality of the data (function: RunPCA). The FindNeighbors function was then used to construct a Shared Nearest Neighbor (SNN) graph on the dimensionally reduced data from the first 10 principal components. For Human Lung dataset, we further removed the effect from technical or biological confounders using Harmony, and SSN graph was constructed using corrected PCA embeddings^21^. The FindClusters function was utilized to identify the cell clusters. Each cluster was identified by differentially expressed genomic signatures or automated cell type annotation. For automated cell type annotation, we used the scCATCH package ^22^ where the findmarkergenes function is applied with *p*-value threshold of 0.05 and logFC threshold of 0.25 to find marker genes for each cluster and then the scCATCH function is utilized to identify the corresponding cell types. To perform DGE analysis between the aged and young cells, we subset the Seurat object by cell type and applied the FindMarkers function with ident.1 = “old” and ident.2 = “young” and the default parameters.

For the bulk RNA-seq data from the human blood, we used the *limma* R-package^23^ to regress out the effect of gender and smoking status of the selected subjects from the COPDGene dataset. The differentially expressed genes were defined as the genes with FDR adjusted *p*-value <0.05 and abs(log2-fold change) >0.25 for all datasets.

### Regulon Activity

The set of transcription factors (TF) and their transcriptional targets (i.e., regulons) used for the analysis of the human and mouse lung datasets were defined using DoRothEA^10,24,25^. Each regulon is considered as a gene set and converted into a GeneSet object using the GeneSet() function from the GSEABase package^26^. We used the Escape package^27^ to calculate Single Sample Gene Set Enrichment (ssGSEA) scores for each regulon using the enritchIt() function. The ssGSEA scores were calculated for the young and old cells in each cell type separately and the differential ssGSEA score was calculated by subtracting the average score of the young cells from the average score of the old cells (i.e., *diff_score = avg(ssGSEA_old) – ssGSEA_young)*). We used the wilcox.test() function in R to calculate the Wilcoxon p-value and used the p.adjust() function to find the adjusted p-values using the false discovery rate method (FDR) for all regulons.

### Cell-cell communication analysis

We used the CellChat R package (version 1.1.3)^28^ to infer and analyze intercellular communication via ligand-receptor integrations. CellChat utilizes the expression of known ligand-receptor pairs in different cell types to infer cell-cell communication. Conserved and context-specific signaling pathways were identified by comparing the overall information flow at different conditions (age group). The information flow of a signaling pathway represents the overall communication probability among all pairs of cell types in the ligand-receptor network. Only signaling pathways with a difference of scaled information flow >10 and P value <0.05 were selected for visualization.

### Cell culture

Primary mouse embryonic fibroblasts (MEF) were isolated from wild-type C57BL/6J mice as previously described and cultured in DMEM (4.5g/L glucose) supplemented with 10% FBS for different days at 37 °C with 5% CO2.

### pBabe-LEF1 construct and retroviral infection

pBabe-puro LEF1 (the long isoform) was a gift from Joan Massague (Addgene plasmid # 27023; http://n2t.net/addgene:27023; RRID:Addgene_27023). Retroviruses were produced by transiently transfecting HEK293T cells with a mixture of pBabe/pBabe-LEF1 and pCL-ampho plasmids. 72 hours after transfection, retroviruses were collected and used to infect day 1MEFs with a MOI of 3 for 48 hours in the presence of 8μg/ml polybrene.

### Immunoblots

Whole cell lysates were isolated from MEFs with RIPA buffer (150Mm NaCl, 1% Triton X-100, 0.5% sodium deoxycholate, 0.1% SDS, and 50mM Tri (pH 7.4) with protease and phosphatase inhibitors). Protein concentrations were determined by Bradford assays (Bio-Rad, USA), and aliquots were snap-frozen in liquid nitrogen and stored at -80°C until usage. Immunoblot analyses were performed as described previously 25. Briefly, 25μg proteins separated by 4∼20% gradient SDS-PAGE gels (Bio-Rad) were transferred to nitrocellulose membranes, blocked in TBST buffer supplemented with 5% bovine serum albumin (BSA) or 5% fat-free milk and incubated overnight with primary anti-LEF1 antibody (Cell Signaling, #2230), anti-p16 antibody (ThermoFisher, # PA5-20379) at 4°C overnight. Blots were incubated with an appropriate secondary antibody coupled to horseradish peroxidase at room temperature for 1 hour and reacted with ECL reagents per the manufacturer’s (Thermo Scientific, USA) suggestion and detected by Biorad ChemiDoc MP Imaging System.

Human lung tissues were homogenized and lysed in RIPA buffer. Proteins were quantified by Pierce™ BCA Protein Assay (Thermo Scientific). Equal amounts of proteins (50μg) from cell preparations were separated by sodium dodecyl sulfate-polyacrylamide gel electrophoresis (SDS-PAGE) and electrotransferred to a PVDF membrane (Bio-Rad) using a Trans-Blot Turbo™ transfer system (Bio-Rad). After transfer, membranes were washed in TBS-T (10 mM Tris, pH 8.0, 150 mM NaCl, 0.05% Tween 20), and blocked with TBS-T supplemented with 5% non-fat dry milk (Dry Powder Milk, RPI, IL, USA) for 1 hour at RT. Then, membranes were incubated overnight at 4°C with different primary antibodies in TBS-T against LEF-1 (1:500, Antibody# 2230, Cell Signaling Technology, MA, USA), and β-actin (1:30000, A3854, Sigma-Aldrich, USA). Next day, membranes were washed, and incubated with horseradish peroxide-conjugated secondary antibody (1:2000, Antibody#7074, Cell Signaling Technology) for 1 hour at RT. Following additional wash steps with TBS-T, membranes were treated with Clarity™ western ECL substrate (Bio-Rad). Quantification was performed by measurement of signal intensity with Image J software (National Institute of Health, Bethesda, MD, USA). Statistical analysis was performed using the pairwise non-parametric Mann-Whitney test from the GraphPad Prism software.

### qRT-PCR

Total mRNA was isolated from MEFs with PureLink RNA mini kit (Life Technologies) with additional on-column DNase digestion step to remove genomic DNA per the manufacturer’s instructions. Reverse transcription was carried out using 5μg of RNA using Superscript III (Life Technologies) per the manufacturer’s instructions. For gene expression analyses, cDNA samples were diluted 1/30-fold (for all other genes except for 18sRNA) and 1/900-fold (for 18sRNA). qPCR was performed using the SYBR green system with sequence-specific primers (**Table S2**, Supplementary Materials). All data were analyzed with 18S or β-actin as the endogenous control and all amplicons span introns. Data were analyzed and presented with GraphPad Prism software. Plots show individual data points and bars at the mean and ± the standard error of the mean (SEM). One-tailed t-tests were used to compare means between groups, with significance set at p<0.05.

### Senescence-associated β-galactosidase (SA-β-gal) assay

Cells were washed twice with phosphate-buffered saline (PBS; pH 7.2), fixed with 0.5% glutaraldehyde in PBS and washed in PBS supplemented with 1 mM MgCl2. Cells were stained at 37°C in X-Gal solution (1 mg/mL X-Gal, 0.12 mM K3Fe[CN]6, 0.12 mM K4Fe[CN]6, 1 mM MgCl2 in PBS at pH 6.0). The staining was performed for 24h at 37°C.

## RESULTS

### Age-related gene expression changes are cell-and species-specific

To uncover important drivers of aging, we examined publicly available human and murine scRNA-seq lung datasets. The datasets included young and aged healthy donors. As a first step, we clustered the different cell types (**Figure 1**, UMAPs and Figure S1) and we identified seven cell types that were found in both datasets (i.e., Macrophages, T Cells, B Cells, AT1, AT2, Club Cells, and Endothelial Cells). The mouse lung dataset included a separate cluster of Dendritic Cells whereas the human lung dataset included separate clusters for Monocyte, Fibroblast, Ciliated, Mast, Lymphatic Endothelial, and Smooth Muscle Cells. Subsequently, we performed Differential Gene Expression (DGE) analysis between samples from old and young donors, focusing on immune cell types that were well represented in both datasets. Further analysis showed that each cell type displayed significant age-related changes in the expression of multiple genes but, only twenty-four genes were found in all immune cell types (**Figure 1**, top volcano plots): NDUFA13, RPS4Y1, HNRNPC, ANKRD28, LRRFIP1, FKBP5, TSC22D3, KLF6, FOS, NBEAL1, CD44*ATP5D, EIF5A, CST3, S100A11, GRN, FIS1, S100A9, SNHG8, NDUFA3, ATP5I, NDUFB1, POLR2L and IFITM3*. Similarly (**Figure 1**, bottom volcano plots), we found eight genes significantly changing with age across Macrophages, T cells, and B cells in the murine dataset: *Scgb1a1, Igkc, Malat1, mt-Rnr2, Gm26924, Ighm, Igha, Igj*. Interestingly, none of the differentially expressed genes was common for both species.

**Figure 1.**
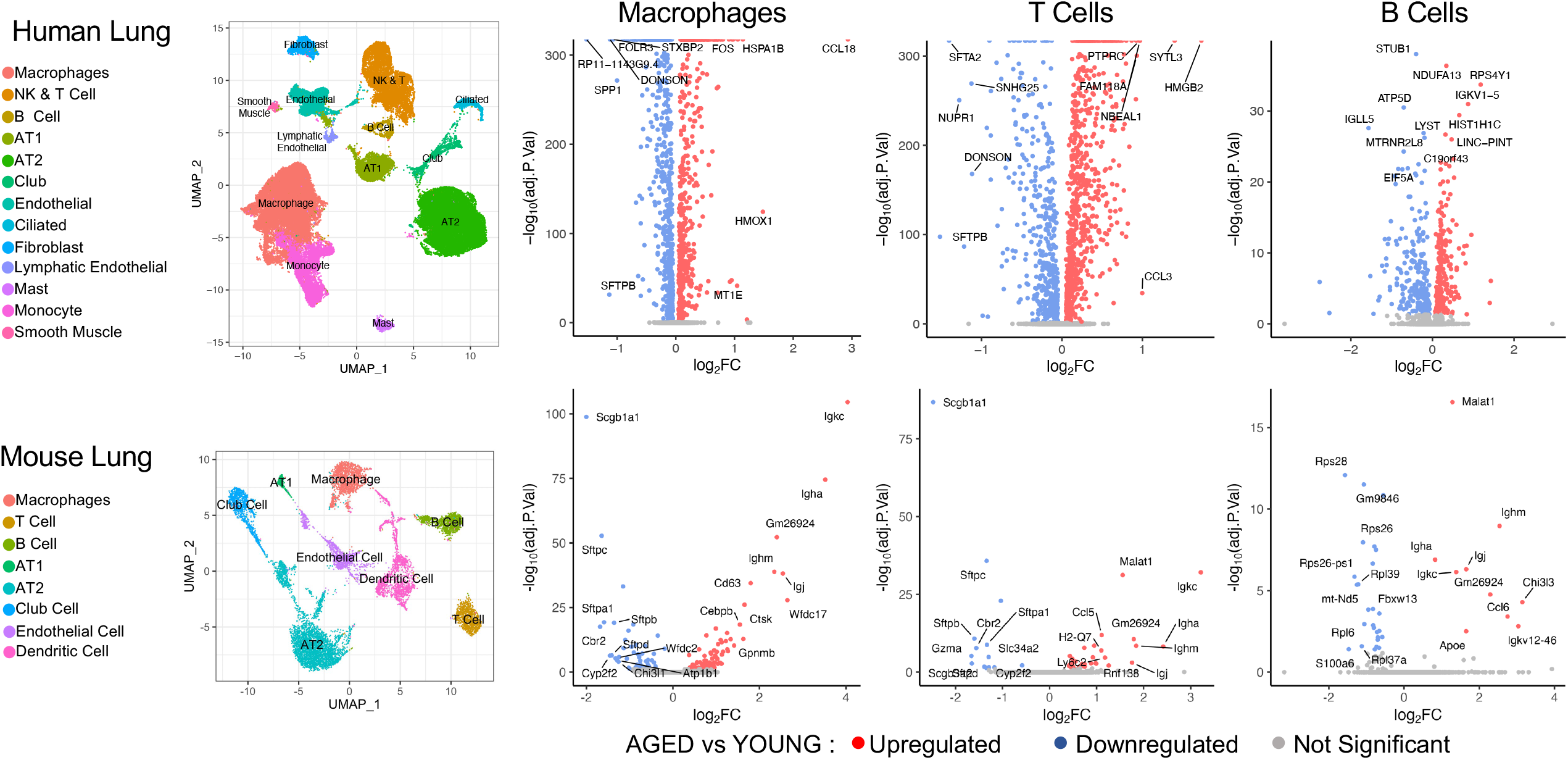
Identification of differentially expressed genes in the human and mouse lung datasets. UMAP plots show the main clusters in the human and mouse lung datasets. Volcano plots show differentially expressed genes in human and murine lung Macrophages, T Cells, and B cells. Significance threshhold is set at adj.P-value<0.05. The names of the top10 highly upregulated/downregulated genes are shown.

### Cell-cell communication analysis revealed dysregulation of age-related cellular pathways

CellChat algorithm^28^ exploits the built-in information of a signaling molecule interaction database and infers a quantitative detection of intercellular communication networks. By comparing the information flow, generated for each signaling pathway that was identified in lung cells from both young and aged donors, we were able to determine age-related changes in several cellular pathways. In aged cells, we observed a dramatic increase in ANXA1, GALECTIN, and IL1 network activity while TENASCIN and SEMA3 networks suffered a significant decrease. To a lesser extent, UGRP1, CLEC, MIF, and ANGPTL network activities were also increased in aged cells while ITGB2 decreased (**Figure 2A**). ANXA1 (Annexin 1) has a known anti-inflammatory role^29,30^ and increased levels could indicate a response to an age-related inflammatory state (inflammaging)^31^. Galectin levels are increased with age^32^ which has been associated with several diseases^33^. Conversely, low levels of galectin are considered a good indicator of successful, healthy aging^34^. Galectin is also anti-inflammatory^35^ and increased levels could suggest a response to inflammaging. Proinflammatory IL1 cytokine signaling has been implicated in inflammation of several tissues including the lung^36^.

**Figure 2:**
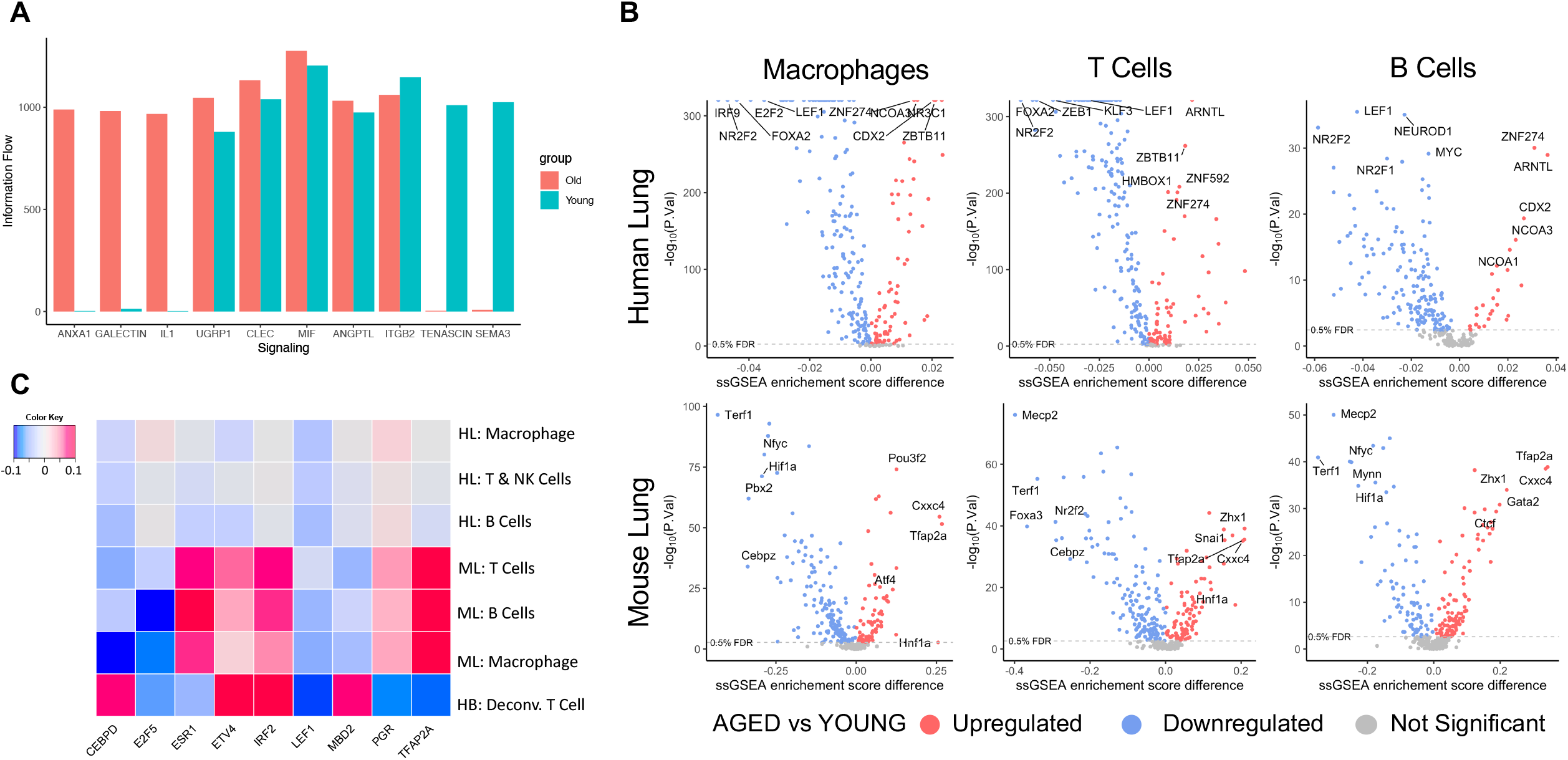
Age-related differences in the activity of cell communication networks and regulons. **A**) Information Flow of CellChat inferred signaling ligands in young and aged human lungs. **B**) Calculation of the differential regulon activity (ssGSEA score difference) in human lung Macrophages, T Cells, B cells (TOP), and murine Macrophages, T Cells, B Cells (BOTTOM). **C**) Identification of common regulons between human (lung and blood) and mouse (lung) cells. Heatmap shows the difference between the average regulon ssGSEA scores in old versus young samples, with FDR <0.1 in all cell types. HL: Human Lung, ML: Mouse Lung, HB: Human Blood.

### LEF1 regulon is consistently found less active in aged subjects

Focusing on the immune cells that were used in our DEG analysis, we compared regulon activity between young and aged donors. To this end, we calculated ssGSEA scores for all regulons included in the DoRothEA gene regulatory network and found that 141 regulons were significantly changed with age in humans (Wilcoxon p value <0.05) (**Figure 2B**, top) while 132 regulons were significantly changed in mice (**Figure 2B**, bottom). Based on differential regulon activity, we identified 70 regulons showing a significant age-related change in both human and murine lung cells. To narrow down the search for factors that could have a more universal role in aging, we analyzed data from the COPDGene study. We calculated the differential ssGSEA scores of the blood samples from old and young healthy donors, compared them to our previous results, and found nine common regulons (CEBPD, E2F5, ESR1, ETV4, IRF2, LEF1, MBD2, PGR, and TFAP2A) that were significantly changed in our human lung, mouse lung, and human blood samples. Of these nine, only *LEF1* regulon activity was consistently decreased with age in all datasets (**Figure 2C**).

### LEF1 expression is dysregulated during cellular senescence

Since cellular senescence is a major hallmark of aging^2^, we next examined whether LEF1 expression also changes during cellular senescence. Primary mouse embryonic fibroblasts undergo spontaneous cellular senescence in culture, characterized by flattened cell morphology, increased expression of senescence marker CDKN2A/p16INK4a/p16, and increased senescence-associated β-galactosidase (SA-β-Gal) activity (**Figure 3A, D**)^37,38^. Interestingly, we detected two LEF1 isoforms that were regulated in a completely anti-parallel fashion. While the longer LEF1 isoform (WT-LEF1) (∼60kD) was decreasing to undetectable levels during cellular senescence, the shorter isoform (∼40Kd) gradually increased (**Figure 3A**). The size of the shorter protein is consistent with the size of a dominant-negative LEF1 isoform, lacking the N-terminus β-catenin-binding domain^39^.

**Figure 3.**
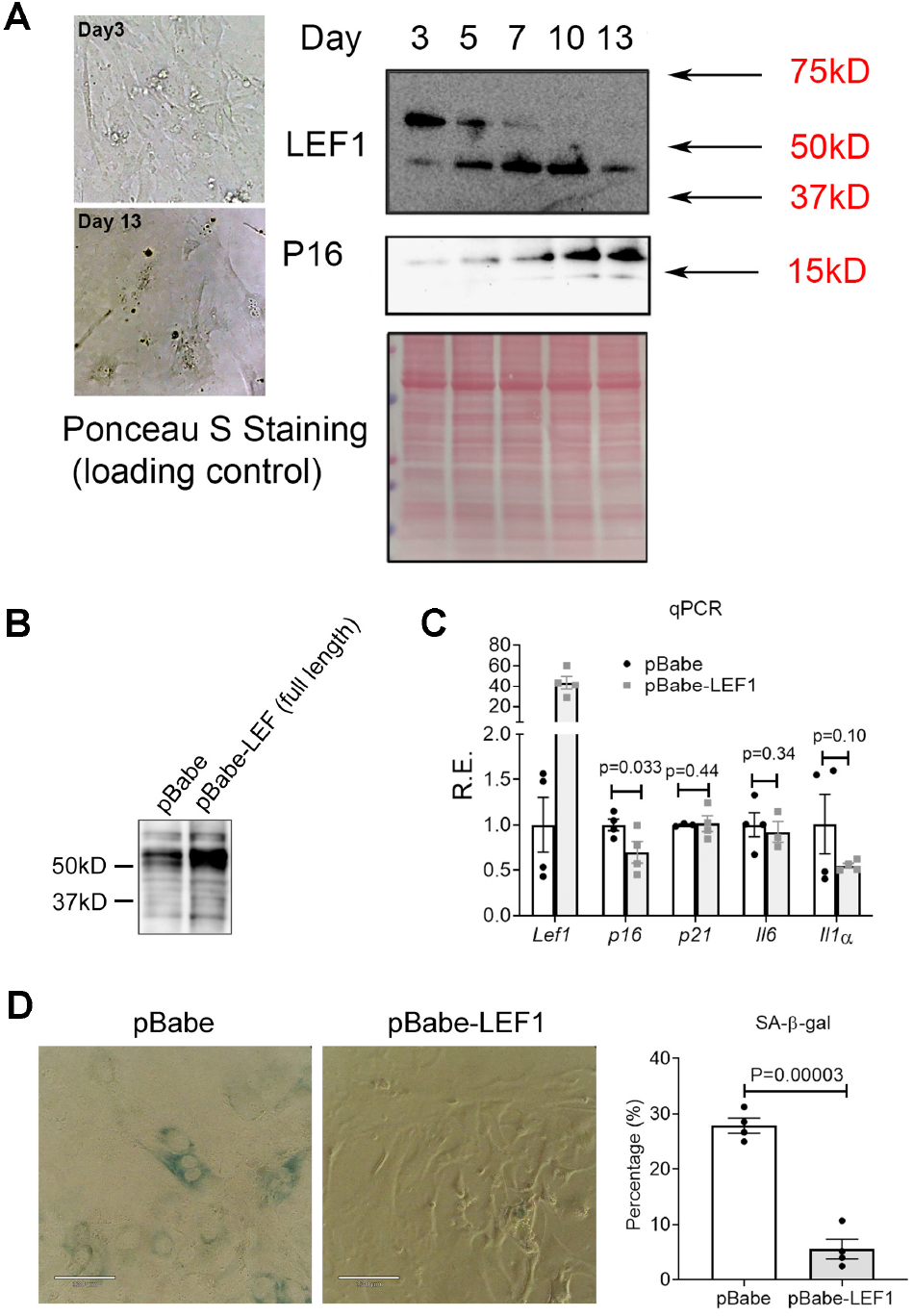
LEF1 is implicated in cellular senescence. **A**) Representative images of the morphology of MEFs at day3 and day13 after culturing. Immunoblot detection of p16 and LEF1 expression at different days. Ponceau S staining is provided as a loading control. **B**) Immunoblot detection of LEF1 in cells expressing pBabe-empty vector or pBabe-LEF1 (long isoform) construct. **C**) qPCR analysis of the expression of *Lef1* and four common senescence markers at day 9 (8 days post retroviral infection) in MEFs. Results are shown as relative expression changes compared to cells transfected with the empty vector. **D**) Representative images (left) and quantification (right) of SA-β-Gal staining in control or LEF1-overexpressing MEFs at day 9 (8 days post retroviral infection).

To investigate whether the decrease in the longer LEF1 isoform contributes to cellular senescence, we ectopically expressed the longer LEF1 isoform in MEFs at day 1 and quantify its impact on cellular senescence progression (**Figure 3B**). The ectopic longer LEF1 isoform expression resulted in the significant downregulation of p16 expression but did not affect some other of the commonly cellular senescence markers, although *Il1α* expression was marginally reduced (**Figure 3C**). Nevertheless, staining with β-galactosidase showed a significant decrease of more than 3-fold, in the number of senescent cells upon the ectopic expression of longer isoform LEF1 (**Figure 3D**). Together, these results demonstrate a possible role of the longer isoform of LEF1 in partially attenuating cellular senescence.

### LEF1 regulon activity in IPF lungs

Due to the important role aging and cellular senescence play in the pathophysiology of Idiopathic Pulmonary Fibrosis (IPF) ^40^, we checked LEF1 expression in lung tissue from Normal/Healthy and IPF lung donors (**Figure 4A, 4B**). Since IPF patients tend to be older and their phenotype is shaped by both age and disease, we included in our analysis aged-matched healthy lung donors. Immunoblot assays of lung tissue showed an overall increase in LEF1 protein in older healthy and IPF donors. Quantification of the two isoforms in all donor groups revealed a higher increase in the short isoform (FC=4.9) than the long isoform (FC= 3.7) in healthy old lungs compared to healthy young lungs. Although less significant, the same trend was observed in IPF lungs where the short and long isoforms showed a 3.4 versus a 2.1 fold change increase respectively. When the relative abundance of both isoforms was evaluated as the ratio to the total LEF1 signal, we observed a significant increase in the relative amount of the short LEF1 isoform (**Figure 4B**) coupled with a significant decrease in the relative amount of the long LEF1 isoform.

**Figure 4.**
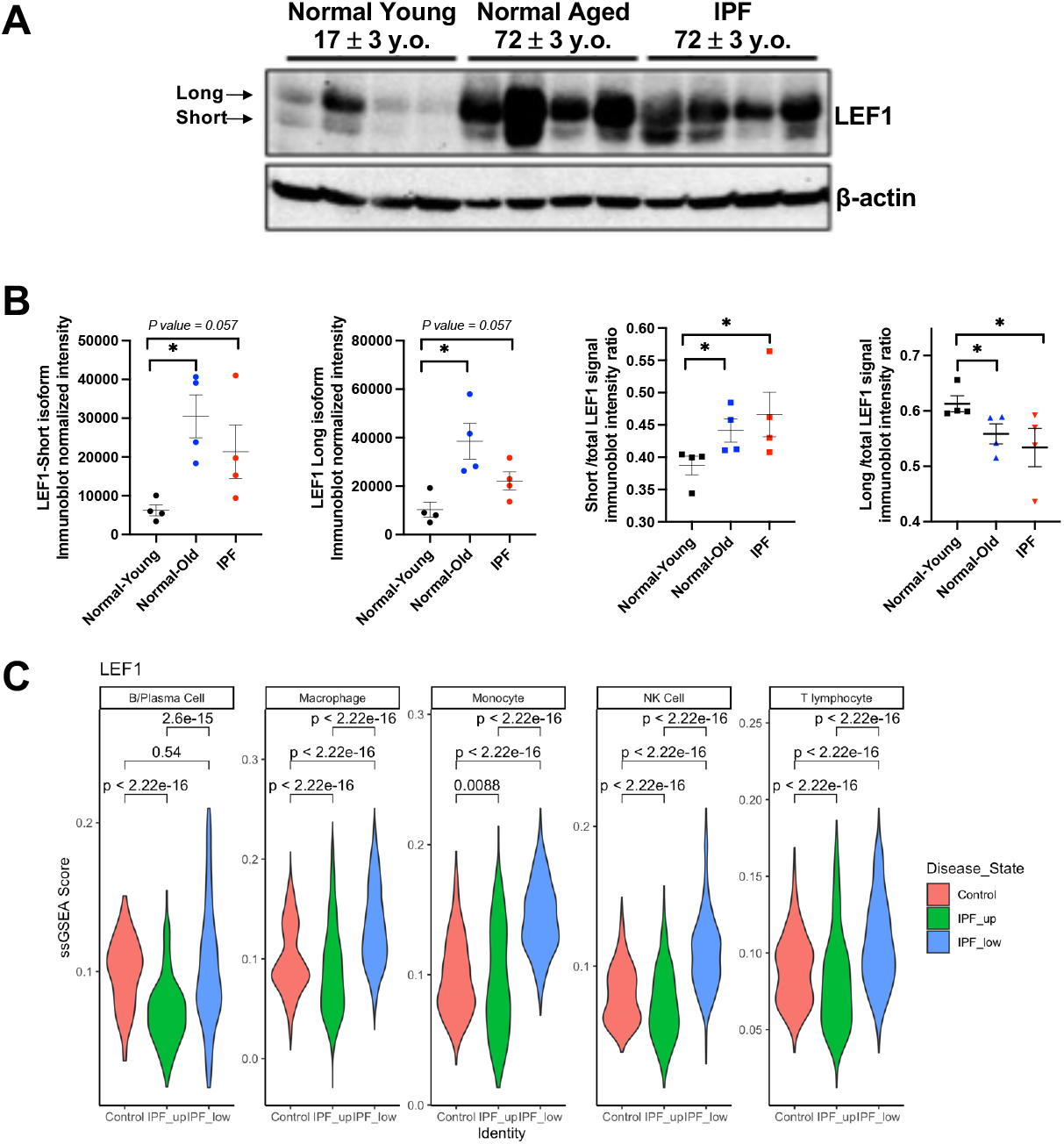
LEF1 expression in human lung tissue lysates. **A**) LEF1 immunoblot of four normal healthy young, four healthy old, and four IPF human lung tissue lysates. **B**) Quantification of LEF1 isoforms using β-actin used as a loading control. Data represent the mean value ± SEM (n = 4 samples evaluated per group). *p <0.03, pairwise non-parametric Mann-Whitney test. **C**) Violin plot graphs showing LEF1 ssGSEA scores in human lung immune cells. IPF: Idiopathic pulmonary fibrosis, IPF-up: upper lung lobe, IPF-low: lower lung lobe.

We have previously shown that IPF lung topology greatly affects the degree of disease severity: upper lung lobes typically appear less scarred and often show very little histological change while lower lung lobes are greatly affected by the disease^11^. We examined the activity of LEF1 regulon in lung immune cells from our data set **(Figure 4C)**, and found that overall, it was decreased in the upper lobe of IPF lungs, supporting the presence of aging/senescent cells. Interestingly, immune cells from the lower and highly fibrotic IPF lung lobes show increased LEF1 regulon activity, suggesting a more complex LEF1 regulatory pathway in advanced fibrosis that is not limited to cellular senescence.

## DISCUSSION

Our study addresses the important topic of mechanisms of aging and aging-related pathologies. The study of such multiparametric phenomena could benefit from computational approaches that take advantage of the increasing number of studies that include samples from a wide spread of age groups. Many of these studies do not directly question the effects of aging, but their control samples form a really underexplored valuable resource. For our study, we analyzed and compared data from three different sources: scRNAseq data from an IPF-related study, blood bulk RNAseq data from healthy smoker population of COPDGene®, and scRNAseq data from an aging-related mouse study. One limitation of our approach is that we aimed to compare results in different cell types across all three datasets, thus our analysis was confined to macrophages, T cells and B cells, which these samples have in common. Taking into consideration the key role of inflammation in aging^31,41^, our results offer a significant contribution to the field.

Our aim was to identify key pathways that are affected throughout aging in all immune cells under investigation. Based on previous studies, we were expecting that the conservation of differentially expressed genes across tissues and species will be limited^5^, but we wanted to focus at a higher level of gene regulation: regulatory modules. Comparing regulon activity in all cell types and species, we observed ubiquitous age-related differences in thirteen regulons. LEF1 regulon was the only regulon with consistent pattern of age-dependent activity in both human and murine immune cells. Notably, although the overall LEF1 regulon activity invariably decreasing with age, the expression profiles of several LEF1 target genes did not change consistently among the various cell types and across organisms, indicating that downstream signals may be diverse and cell-type specific. This could explain why we and other studies were unable to detect significant matches in DGE profiles of the immune cells; and it is consistent with previous reports that aging could have a subtle transcriptional signature depending on the tissue type^5^, complicating the straightforward detection of ubiquitous aging-driving factors.

Immunodetection of LEF1 in Mouse Embryonic Fibroblasts revealed a senescence-associated dysregulation of two distinct isoforms. The longer isoform showed a steady decrease with increased replicative senescence (indicated by increase in p16, a known senescence marker^38^), while the short isoform was gradually upregulated. Ectopic expression of the longer isoform prevented replicative senescence, supporting the role of the full-length LEF1 in cell proliferation^42^. The two LEF1 isoforms were also detected in tissue lung samples from healthy and IPF donors. It is important to note that lung tissue in general and IPF fibrotic tissue in particular present a challenging material for analysis due to the high cell type heterogeneity and the spatial diversity of the disease progression. Whereas lung immune cells from older subjects show a less active LEF1 regulon, suggesting a possible decrease in the long LEF1 isoform expression, protein levels were increased in both healthy and IPF lung tissue. Interestingly, in old and IPF lungs, the relative amount of the long isoform decreased while the short isoform increased, which is reminiscent of the LEF1 dysregulation pattern in our senescent MEFs. Unsurprisingly, the LEF1 regulon activity was decreased in the upper lung lobes of IPF patients in agreement with our findings in old healthy lungs. This suggests that immune cells from older lungs as well as normal-looking areas from IPF lungs could undergo a LEF1/senescence-related decline. LEF1 regulon activity in the immune cells of the highly fibrotic lower lobes appears elevated, suggesting a different and rather more proliferative state in these cells.

LEF1 is a transcription factor expressed in B and T cells^43,44^. It is implicated in various cancers where its overexpression is associated with poor prognosis^45,46^. LEF1 interacts with beta-catenin^47^ and SMAD2-3^48^, key molecules in the Wnt and TGF-β pathways respectively. Interestingly, Wnt and TGF-β signaling pathways can independently or synergistically regulate LEF1 targets^48^. Both pathways are dysregulated in cellular senescence and aging^49,50^ and LEF1 is found to be downregulated in certain aged tissues ^5^. While the function of long LEF1 isoform and the significance of LEF1 expression levels have been studied, the fluctuation of the smaller isoform, lacking the β-catenin binding domain, has less clear consequences. Some studies suggest that the small isoform acts as a dominant negative variant inhibiting β-catenin binding and consequently promoter activation^39^, while other studies support a more complex role where the short isoform has distinct targets and biological functions^51^.

## CONCLUSIONS

The alternative LEF1 isoform expression is a key regulator of β-catenin pathway with the long isoform promoting and the short isoform inhibiting the expression of the LEF1 target genes (**Figure 5**). We found that functional LEF1 and LEF1 regulon activity is reduced with age in multiple immune cell types from different human and murine tissues. We also showed that two LEF1 isoforms are severely dysregulated in aged lungs. Furthermore, we have provided experimental evidence that LEF1, through its two protein isoforms, is an important regulator of cellular senescence, with the long isoform preventing senescence in cell culture. Without a doubt, more research is required to determine whether LEF1 dysregulation is intrinsic to the aging process, leading to increased vulnerability to cellular malfunction and disease. Clearly though, LEF1 dysregulation, and by extension, the dysregulation of its downstream cellular factors and pathways, appears to be cell-specific, which dictates more targeted and precise approaches for future aging studies.

**Figure 5.**
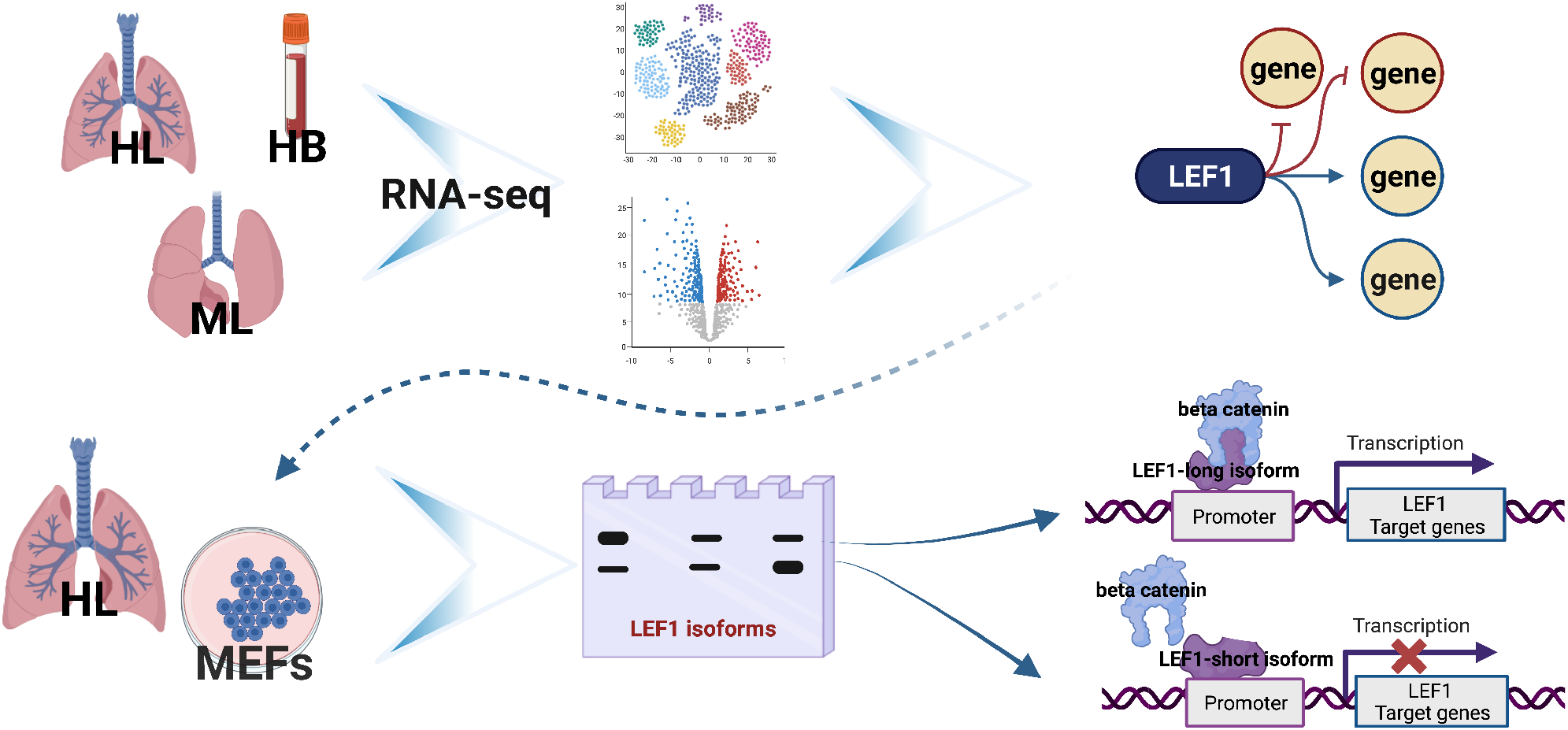
Graphical summary of the findings of the current study. HL: Human lung, ML: murine lung, HB: Human Blood, MEF: mouse embryonic fibroblasts.

## DECLARATIONS

### Ethics and approval consent to participate

Not applicable.

### Consent for publication

Not applicable.

### Availability of data

Data were retrieved from public databases (GEO <u>GSE128033</u>, <u>GSE122960</u>, <u>GSE124872</u>, <u>GSE158699</u>,

### Authors contributions

PVB and MGK designed the study with the help of MJ and KS. MJ, KS performed all computational analyses. WD, LR, SI performed the experiments under the supervision of MR, BZ. MJ, KS, MGK, PVB wrote the paper with input from ALM, RAL, MR, BZ. All authors have read and approved this manuscript.

## FIGURE LEGENDS

***Supplementary Figure S1***. UMAP plot visualizes human dataset. (A) cells colored by age/young state (B) cells colored by sample

***Supplementary Figure S2***. UMAP plots showing macrophage cells from GSE122960. Donor-4 (red box) shows an abnormal cell distribution compared to the other donors.

***Supplementary Figure S3***. Lung histopathology of samples from the two public human scRNA-seq datasets. Donor-2 (red box) histopathology is indicative of smoking or pollution exposure.

## REFERENCES

1. Lauren Medina; Shannon Sabo; Jonathan Vespa; U.S. Census Bureau. Living longer : historical and projected life expectancy in the United States, 1960 to 2060. U.S. Census Bureau, Current population reports.2020:no. 1145.

2. López-Otín C, Blasco MA, Partridge L, Serrano M, Kroemer G. Hallmarks of aging: An expanding universe. Cell 2023;186:243–78.

3. Di Micco R, Krizhanovsky V, Baker D, d’Adda di Fagagna F. Cellular senescence in ageing: from mechanisms to therapeutic opportunities. Nat Rev Mol Cell Biol 2021;22:75–95.

4. Rodier F, Campisi J. Four faces of cellular senescence. J Cell Biol 2011;192:547–56.

5. Barth E, Srivastava A, Stojiljkovic M, et al. Conserved aging-related signatures of senescence and inflammation in different tissues and species. Aging (Albany NY) 2019;11:8556–72.

6. Franks TJ, Colby TV, Travis WD, et al. Resident cellular components of the human lung: current knowledge and goals for research on cell phenotyping and function. Proc Am Thorac Soc 2008;5:763–6.

7. MacNee W. Is Chronic Obstructive Pulmonary Disease an Accelerated Aging Disease? Annals of the American Thoracic Society 2016;13:S429–S37.

8. Mora AL, Bueno M, Rojas M. Mitochondria in the spotlight of aging and idiopathic pulmonary fibrosis. The Journal of Clinical Investigation 2017;127:405–14.

9. Cho SJ, Stout-Delgado HW. Aging and Lung Disease. Annual Review of Physiology 2020;82:433–59.

10. Holland CH, Szalai B, Saez-Rodriguez J. Transfer of regulatory knowledge from human to mouse for functional genomics analysis. Biochimica et Biophysica Acta (BBA)-Gene Regulatory Mechanisms 2020;1863:194431.

11. Morse C, Tabib T, Sembrat J, et al. Proliferating SPP1/MERTK-expressing macrophages in idiopathic pulmonary fibrosis. European Respiratory Journal 2019;54:1802441.

12. Cruz T, Jia M, Sembrat J, et al. Reduced Proportion and Activity of Natural Killer Cells in the Lung of Patients with Idiopathic Pulmonary Fibrosis. American journal of respiratory and critical care medicine 2021;204:608–10.

13. Reyfman PA, Walter JM, Joshi N, et al. Single-Cell Transcriptomic Analysis of Human Lung Provides Insights into the Pathobiology of Pulmonary Fibrosis. Am J Respir Crit Care Med 2019;199:1517–36.

14. Angelidis I, Simon LM, Fernandez IE, et al. An atlas of the aging lung mapped by single cell transcriptomics and deep tissue proteomics. Nature communications 2019;10:1–17.

15. Regan EA, Hokanson JE, Murphy JR, et al. Genetic epidemiology of COPD (COPDGene) study design. COPD: Journal of Chronic Obstructive Pulmonary Disease 2011;7:32–43.

16. Stuart T, Butler A, Hoffman P, et al. Comprehensive integration of single-cell data. Cell 2019;177:1888-902. e21.

17. Newman AM, Steen CB, Liu CL, et al. Determining cell type abundance and expression from bulk tissues with digital cytometry. Nature biotechnology 2019;37:773–82.

18. Lun AT, Riesenfeld S, Andrews T, Gomes T, Marioni JC. EmptyDrops: distinguishing cells from empty droplets in droplet-based single-cell RNA sequencing data. Genome biology 2019;20:1–9.

19. Wolock SL, Lopez R, Klein AM. Scrublet: computational identification of cell doublets in single-cell transcriptomic data. Cell systems 2019;8:281-91. e9.

20. Huang M, Wang J, Torre E, et al. SAVER: gene expression recovery for single-cell RNA sequencing. Nature methods 2018;15:539–42.

21. Korsunsky I, Millard N, Fan J, et al. Fast, sensitive and accurate integration of single-cell data with Harmony. Nature methods 2019;16:1289–96.

22. Shao X, Liao J, Lu X, Xue R, Ai N, Fan X. scCATCH: automatic annotation on cell types of clusters from single-cell RNA sequencing data. IScience 2020;23:100882.

23. Ritchie ME, Phipson B, Wu D, et al. limma powers differential expression analyses for RNA-sequencing and microarray studies. Nucleic acids research 2015;43:e47–e.

24. Garcia-Alonso L, Holland CH, Ibrahim MM, Turei D, Saez-Rodriguez J. Benchmark and integration of resources for the estimation of human transcription factor activities. Genome research 2019;29:1363–75.

25. Holland CH, Tanevski J, Perales-Patón J, et al. Robustness and applicability of transcription factor and pathway analysis tools on single-cell RNA-seq data. Genome biology 2020;21:1–19.

26. Morgan M, Falcon S, Gentleman R. GSEABase: Gene set enrichment data structures and methods. R package version 1540 2021;1.

27. Borcherding N, Vishwakarma A, Voigt AP, et al. Mapping the immune environment in clear cell renal carcinoma by single-cell genomics. Communications biology 2021;4:1–11.

28. Jin S, Guerrero-Juarez CF, Zhang L, et al. Inference and analysis of cell-cell communication using CellChat. Nature Communications 2021;12:1088.

29. Jia Y, Morand EF, Song W, Cheng Q, Stewart A, Yang YH. Regulation of lung fibroblast activation by annexin A1. J Cell Physiol 2013;228:476–84.

30. Rubinstein MR, Baik JE, Lagana SM, et al. Fusobacterium nucleatum promotes colorectal cancer by inducing Wnt/beta-catenin modulator Annexin A1. EMBO Rep 2019;20.

31. Franceschi C, Garagnani P, Parini P, Giuliani C, Santoro A. Inflammaging: a new immune–metabolic viewpoint for age-related diseases. Nature Reviews Endocrinology 2018;14:576–90.

32. de Boer RA, van Veldhuisen DJ, Gansevoort RT, et al. The fibrosis marker galectin-3 and outcome in the general population. J Intern Med 2012;272:55–64.

33. Dong R, Zhang M, Hu Q, et al. Galectin-3 as a novel biomarker for disease diagnosis and a target for therapy (Review). Int J Mol Med 2018;41:599–614.

34. Sanchis-Gomar F, Santos-Lozano A, Pareja-Galeano H, et al. Galectin-3, osteopontin and successful aging. Clin Chem Lab Med 2016;54:873–7.

35. Camby I, Le Mercier M, Lefranc F, Kiss R. Galectin-1: a small protein with major functions. Glycobiology 2006;16:137R–57R.

36. Borthwick LA. The IL-1 cytokine family and its role in inflammation and fibrosis in the lung. Semin Immunopathol 2016;38:517–34.

37. Manning JA, Kumar S. A potential role for NEDD1 and the centrosome in senescence of mouse embryonic fibroblasts. Cell Death Dis 2010;1:e35.

38. Guan Y, Zhang C, Lyu G, et al. Senescence-activated enhancer landscape orchestrates the senescence-associated secretory phenotype in murine fibroblasts. Nucleic Acids Research 2020;48:10909–23.

39. Hovanes K, Li TW, Munguia JE, et al. Beta-catenin-sensitive isoforms of lymphoid enhancer factor-1 are selectively expressed in colon cancer. Nat Genet 2001;28:53–7.

40. Kellogg DL, Kellogg DL, Jr., Musi N, Nambiar AM. Cellular Senescence in Idiopathic Pulmonary Fibrosis. Curr Mol Biol Rep 2021;7:31–40.

41. Fulop T, Witkowski JM, Olivieri F, Larbi A. The integration of inflammaging in age-related diseases. Semin Immunol 2018;40:17–35.

42. Hao Y-H, Lafita-Navarro MC, Zacharias L, et al. Induction of LEF1 by MYC activates the WNT pathway and maintains cell proliferation. Cell Communication and Signaling 2019;17:129.

43. Milatovich A, Travis A, Grosschedl R, Francke U. Gene for lymphoid enhancer-binding factor 1 (LEF1) mapped to human chromosome 4 (q23-q25) and mouse chromosome 3 near Egf. Genomics 1991;11:1040–8.

44. Elyahu Y, Hekselman I, Eizenberg-Magar I, et al. Aging promotes reorganization of the CD4 T cell landscape toward extreme regulatory and effector phenotypes. Sci Adv 2019;5:eaaw8330.

45. Erdfelder F, Hertweck M, Filipovich A, Uhrmacher S, Kreuzer KA. High lymphoid enhancer-binding factor-1 expression is associated with disease progression and poor prognosis in chronic lymphocytic leukemia. Hematol Rep 2010;2:e3.

46. Eskandari E, Mahjoubi F, Motalebzadeh J. An integrated study on TFs and miRNAs in colorectal cancer metastasis and evaluation of three co-regulated candidate genes as prognostic markers. Gene 2018;679:150–9.

47. Behrens J, von Kries JP, Kühl M, et al. Functional interaction of beta-catenin with the transcription factor LEF-1. Nature 1996;382:638–42.

48. Labbé E, Letamendia A, Attisano L. Association of Smads with lymphoid enhancer binding factor 1/T cell-specific factor mediates cooperative signaling by the transforming growth factorbeta and wnt pathways. Proc Natl Acad Sci U S A 2000;97:8358–63.

49. Hu H-H, Cao G, Wu X-Q, Vaziri ND, Zhao Y-Y. Wnt signaling pathway in aging-related tissue fibrosis and therapies. Ageing Research Reviews 2020;60:101063.

50. Tominaga K, Suzuki HI. TGF-β Signaling in Cellular Senescence and Aging-Related Pathology. Int J Mol Sci 2019;20.

51. Edmaier KE, Stahnke K, Vegi NM, Mulaw MA, Buske C. The Short Lef-1 Isoform, Lacking The β-Catenin Binding Domain, Is Not Acting As a Dominant Negative Lef-1 Variant, But Is a Hematopoietic Active Protein With Unique DNA Binding Properties. Blood 2013;122:2415-.

