## Supplemental Figure for "LEF1 isoforms regulate cellular senescence and aging"

### LEF1 alternative transcription regulation affects cellular senescence and aging

#### SUPPLEMENTARY FIGURES

#### Supplementary Figure S1

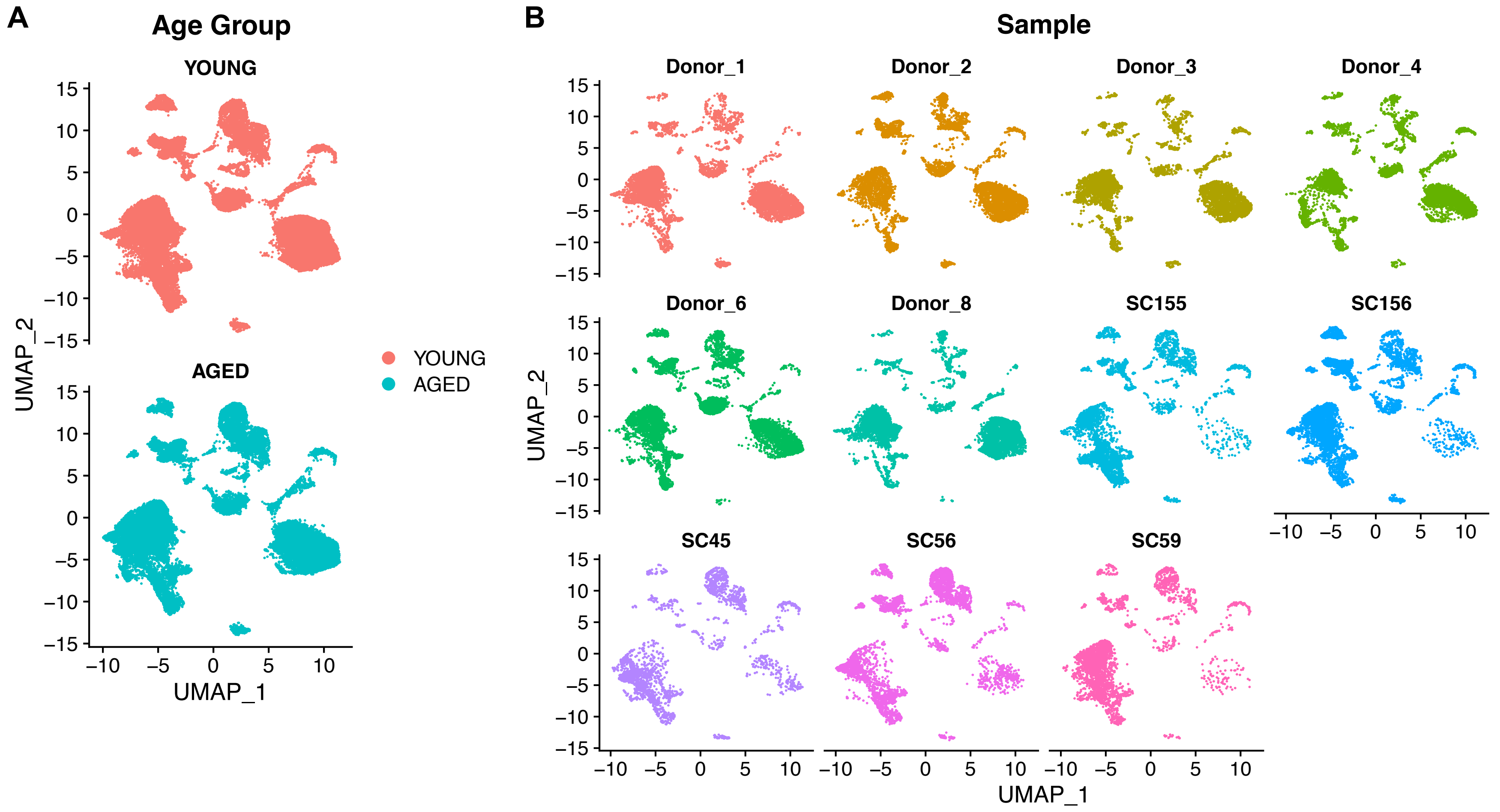

**Supplementary Figure S2**

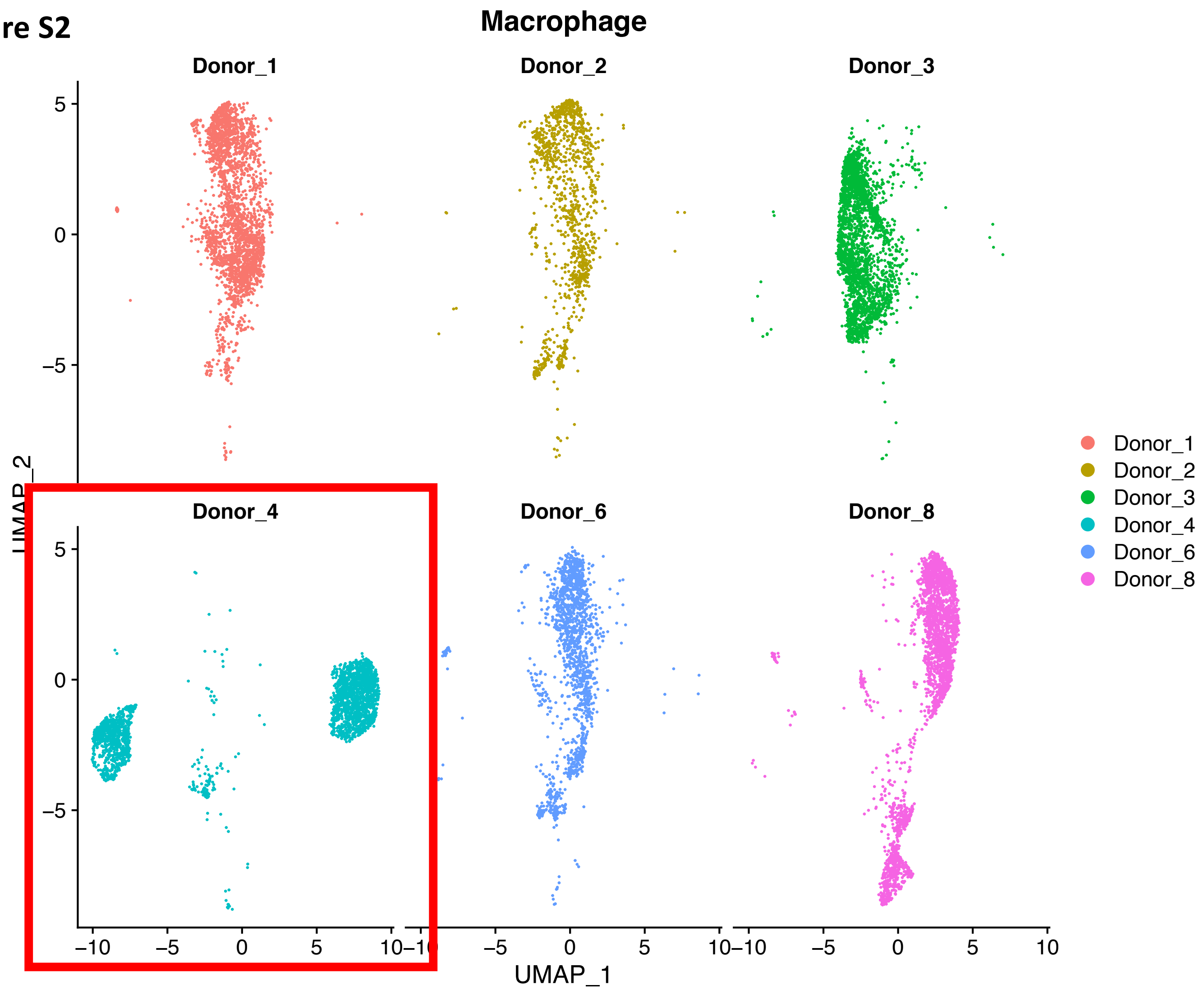

**Supplementary Figure S3**

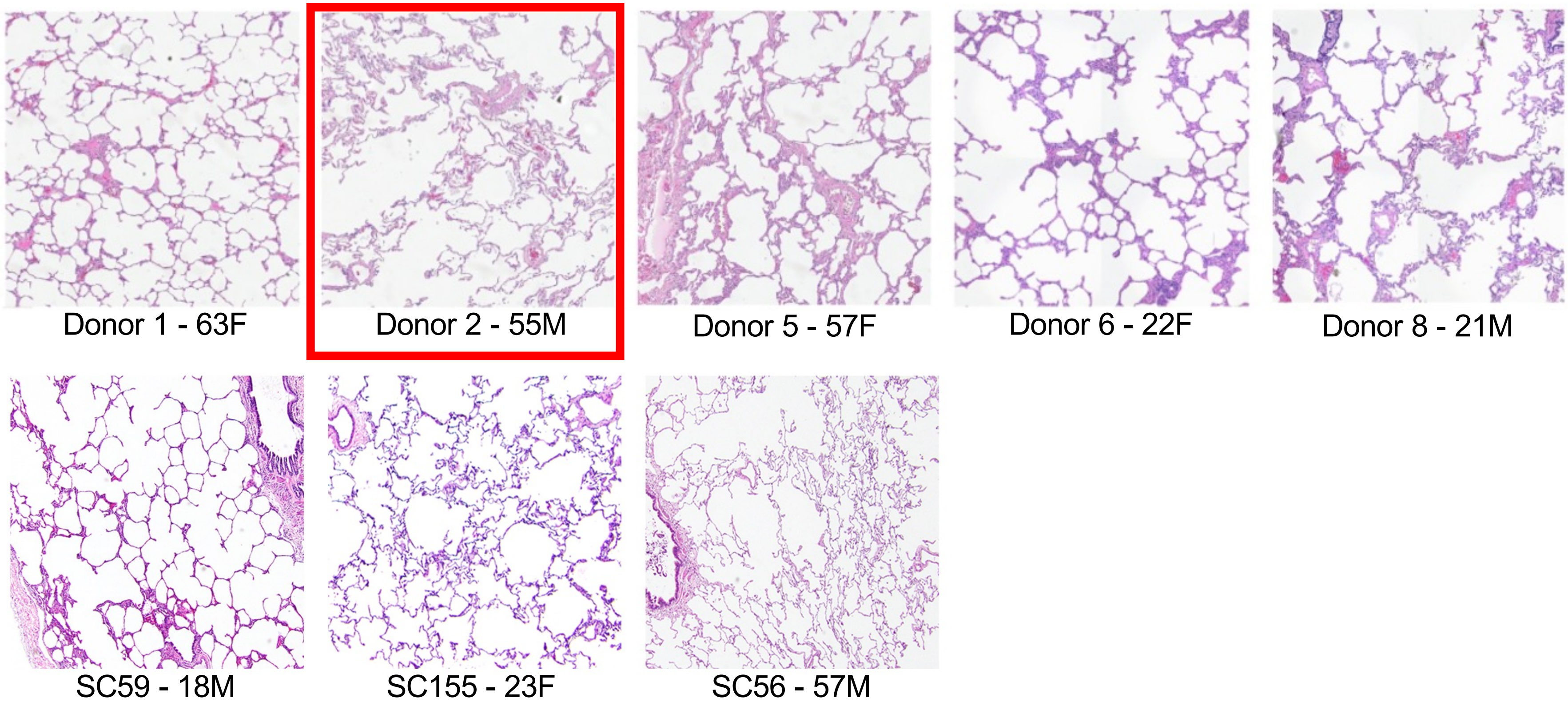
